# Biphasic Modulation of Absence Seizures by Lisuride in GAERS: Dissecting the Role of 5-HT2A Receptors

**DOI:** 10.64898/2026.09.28.755018

**Authors:** Marcello Venzi, Vincenzo Crunelli, Giuseppe Di Giovanni

## Abstract

**Background:** Serotonergic signaling plays an important role in the modulation of absence seizures (ASs), and activation of serotonin 2A receptors (5-HT2ARs) with TCB-2 suppresses spontaneous absence seizures (ASs) in Genetic Absence Epilepsy Rats from Strasbourg (GAERS). Lisuride is a pharmacologically complex serotonergic and dopaminergic ligand with low 5-HT2AR signaling efficacy and a receptor profile distinct from that of TCB-2. We therefore investigated its effects on spontaneous ASs and the contribution of 5-HT2ARs to its actions.

**Methods:** The effects of systemic lisuride (0.1 and 0.5 mg/kg) on spontaneous ASs were examined in freely moving GAERS over 120 min by quantifying time spent in seizure, mean seizure duration and seizure number. The contribution of 5-HT2ARs was investigated by administering lisuride (0.5 mg/kg) following pretreatment with the selective 5-HT2AR antagonist MDL11,939 (0.5 mg/kg).

**Results:** Lisuride produced marked dose- and time-dependent effects on ASs. At 0.1 mg/kg, lisuride elicited modest pro-absence effect. In contrast, 0.5 mg/kg produced a pronounced biphasic response, with a marked reduction in seizure activity during the first 40 min, followed by rebound increase during the second hour. The early suppression involved reductions in both seizure number and duration, whereas the delayed pro-absence effect was predominantly associated with prolonged ASs. MDL11,939 did not prevent the early seizure suppression but blocked the increase in seizure duration while producing a transient increase in seizure number.

**Conclusions:** Lisuride is a dose- and time-dependent bidirectional modulator of ASs in GAERS. The persistence of the profound early anti-absence effect during 5-HT2AR blockade indicates that this response does not require 5-HT2AR activation, whereas the modification of the delayed response by MDL11,939 supports a 5-HT2AR-dependent component in the later phase. These findings distinguish lisuride pharmacologically from TCB-2 and suggest that its biphasic effects arise from the dynamic engagement of multiple receptor mechanisms, potentially including dopaminergic signaling.

## 1. Introduction

Typical absence seizures (ASs) are generalized non-convulsive seizures characterized by transient impairment of consciousness and highly synchronized spike-and-wave discharges (SWDs). Their generation depends on abnormal interactions within cortico-thalamo-cortical networks involving the cerebral cortex, thalamocortical (TC) neurons and the thalamic reticular nucleus [1, 2]. Genetic Absence Epilepsy Rats from Strasbourg (GAERS) reproduce the principal electrographic, behavioral and pharmacological features of human absence epilepsy and provide a well-established model for investigating the mechanisms controlling ASs. [3].

Serotonin (5-HT) is an important modulator of thalamocortical excitability [4], and recent evidence indicates that several 5-HT receptor subtypes influence AS expression. Activation of 5-HT2C receptors (5-HT2CRs) with the agonist Ro 60-0175 suppresses spontaneous ASs in GAERS and reduces the enhanced tonic GABAA inhibition of ventrobasal (VB) TC neurons [5], further supporting the potential cellular link between serotonergic modulation of thalamic inhibition and seizure control [6, 7]. This is consistent with the anti-absence effect of 5-HT2AR activation, as systemic administration of the potent 5-HT2AR agonist TCB-2 markedly reduces ASs in GAERS, whereas the selective antagonist MDL11,939 transiently increases seizure activity (Venzi et al., 2016). Moreover, thalamic 5-HT2AR activation reduces tonic GABAA inhibition in TC neurons and suppresses ASs [8, 9]. Notably, however, 5-HT2ARs can exert opposite effects on thalamic GABAergic signaling in non-epileptic animals, enhancing tonic GABAA inhibition and promoting ASs in Wistar rats, further supporting the context-dependent regulation of thalamocortical excitability by 5-HT2ARs [10].

The consequences of 5-HT2AR activation, however, may also depend on receptor occupancy, intrinsic efficacy, and downstream signaling, and therefore differ considerably among agonists [11-13]. This is particularly relevant because classical serotonergic psychedelics exert their characteristic behavioral effects through 5-HT2ARs, yet not all 5-HT2AR agonists are psychedelic, with lisuride providing a particular example in this context [14, 15].

Importantly, lisuride is not a selective 5-HT2AR ligand but has a broad pharmacological profile involving serotonergic and dopaminergic receptors [16]. Recent work has highlighted a potent 5-HT1A receptor agonism, which contributes substantially to its acute behavioral effects in mice [14]. Despite its multi-receptor profile, lisuride is particularly informative for determining whether the anti-absence phenotype produced by TCB-2 can be reproduced by a pharmacologically distinct, non-psychedelic 5-HT2AR ligand and whether this effect requires 5-HT2AR signaling.

We therefore investigated the effects of lisuride on spontaneous ASs in freely moving GAERS. Lisuride produced a dose- and time-dependent biphasic response, predominantly affecting seizure duration, with early seizure suppression followed by a delayed increase in seizure activity. MDL11,939 did not affect the early anti-absence effect but reversed the delayed increase in seizure burden and duration, while transiently increasing seizure number, indicating distinct mechanisms underlying the biphasic response.

## 2. Materials and Methods

## 2.1 Animals

Adult male GAERS (>P90) were bred from the colony at Cardiff University. Animals were provided with standard diet and water ad libitum and maintained on a 12:12-h light– dark cycle (lights on at 07:00). All procedures were performed in accordance with the UK Animals (Scientific Procedures) Act (1986), ARRIVE guidelines, Basel declaration including the 3R concept, the European Directive 2010/63/EU, and local ethical requirements. All efforts were made to minimize animal suffering and the number of animals used [17].

### 2.2 EEG recordings

Anesthesia was induced with 4% isoflurane in O□ and maintained at 2–3% during stereotaxic surgery. Rats received meloxicam (0.5 mg/kg) postoperatively and were allowed to recover for at least 7 days before EEG recordings were performed in freely moving animals. Animals were habituated to the recording environment before pharmacological experiments [5, 8, 18, 19].

### 2.3 Pharmacological treatments

Following a 1-h habituation period, a stable 1-h baseline EEG and video recordings were obtained before drug administration, followed by continuous EEG and video monitoring for 120 min after treatment. Lisuride was administered systemically at 0.1 or 0.5 mg/kg. These doses were selected with reference to the range previously investigated in GAERS [20].

To investigate the involvement of 5-HT2ARs, the effect of lisuride (0.5 mg/kg) was examined following a 10-min pretreatment with the selective 5-HT2AR antagonist MDL11,939 (0.5 mg/kg). Appropriate vehicle-treated animals were included according to the same experimental design. In all experiments, animals were randomly assigned to the different treatment groups.

### 2.4 Analysis of absence seizures

Spontaneous SWDs were identified according to established electrographic criteria for GAERS [19], and classified as ASs following inspection of the video recordings to ensure that the SWDs in the EEG were accompanied by behavioral arrest.

Seizure activity was quantified using three complementary parameters: total time spent in seizure, average seizure length, and seizure number. Data were analyzed in consecutive 20-min intervals in the control period and throughout the 120-min post-treatment period and normalized to the corresponding baseline where appropriate. This temporal analysis allowed the identification of time-dependent effects that might be obscured by averaging seizure activity across the entire post-treatment period.

### 2.5 Statistical analysis

The effects of systemic administration of the compounds were analyzed using GraphPad Prism version 7 (GraphPad Software, San Diego, CA, USA). Data were analyzed by two-way ANOVA, with drug treatment and time as factors. Dunnett’s multiple-comparisons test was used, where appropriate, to compare each drug treatment with vehicle for the main treatment effect over the full treatment period and for effects at individual 20-min time bins. Time spent in seizures, mean seizure duration, and seizure number were analyzed separately. Data are presented as mean ± SEM. Statistical significance was set at *p* < 0.05. Full statistical results are provided in Supplementary Table 1.

## 3. Results

### 3.1 Lisuride produces dose- and time-dependent modulation of ASs

Systemic administration of lisuride produced dose-, time and time x treatment-dependent effects on spontaneous ASs in GAERS compared to controls (two-way ANOVA; Suppl. Table 1A, *n* = 9 animals per group). At 0.1 mg/kg i.p., lisuride produced a significant overall increase in time spent in seizure (31.9%, *p* < 0.05) and average seizure length (26.9%, *p* < 0.05), without significantly affecting seizure number (Fig. 1A-C). At individual time points, time spent in seizure was unchanged, whereas average seizure length was significantly increased at 80 min (90.7%, *p* < 0.01).

**Figure 1.**
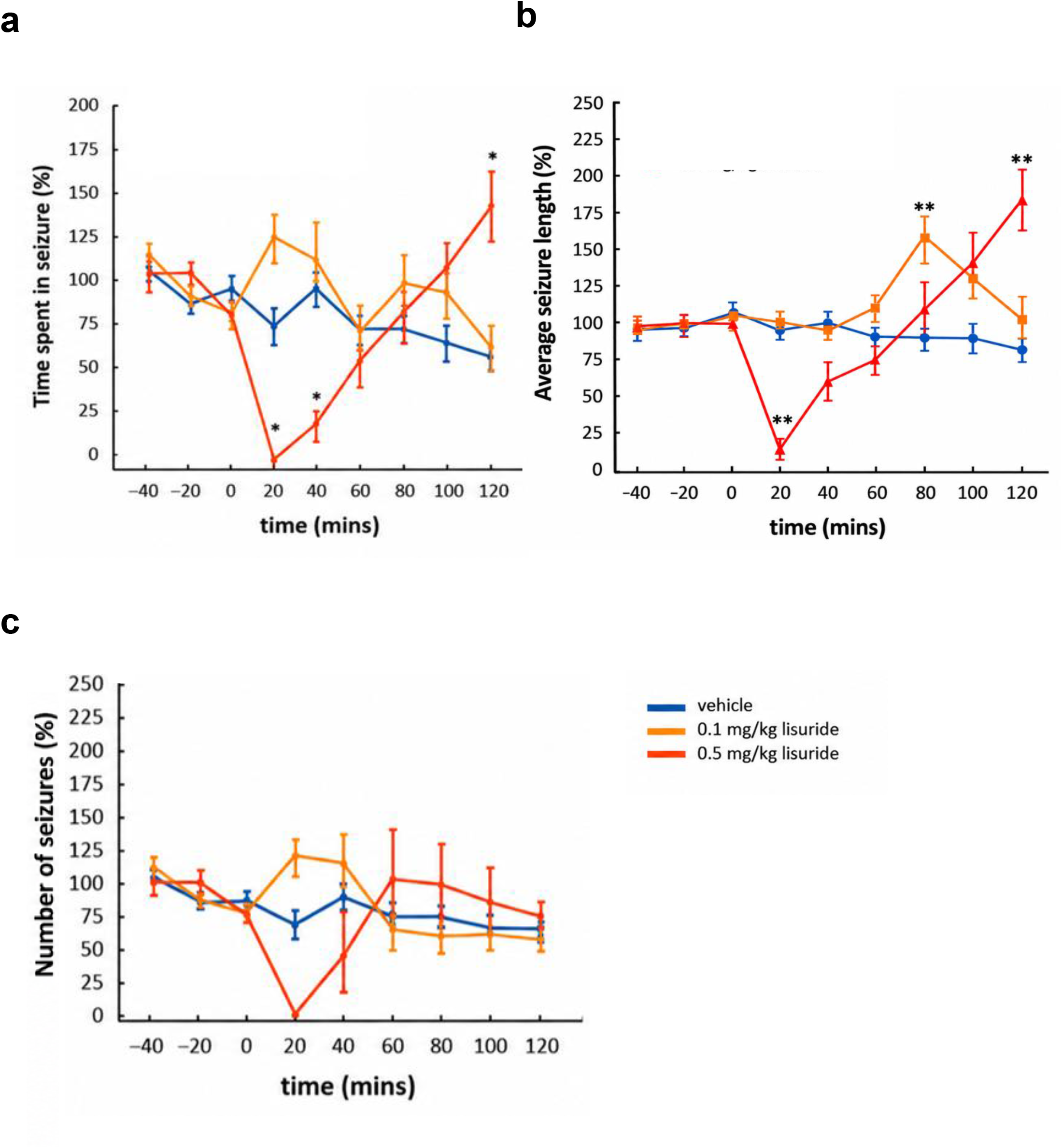
Lisuride produces dose- and time-dependent biphasic modulation of ASs. **(a–c)** Time course of spontaneous absence seizure (AS) parameters following systemic administration of vehicle or lisuride (0.1 and 0.5 mg/kg, n=9 per group). **a**, time spent in seizures; **b**, mean seizure duration; **c**, number of seizures. At 0.1 mg/kg, lisuride produced modest effects, including a significant increase in seizure duration at 80 min. In contrast, 0.5 mg/kg lisuride produced a pronounced biphasic response, characterized by an early suppression of ASs followed by delayed facilitation, predominantly associated with an increase in seizure duration. Data are expressed as a percentage of baseline and presented as mean ± SEM. Asterisks indicate significant differences at the indicated time points: \**p < 0*.*05* and *\*\*p < 0*.*01*. Full statistical results from the two-way ANOVA, including the main effects of treatment and time, their interaction, and Dunnett’s post hoc comparisons are provided in Suppl. Table 1A.

**Figure 2.**
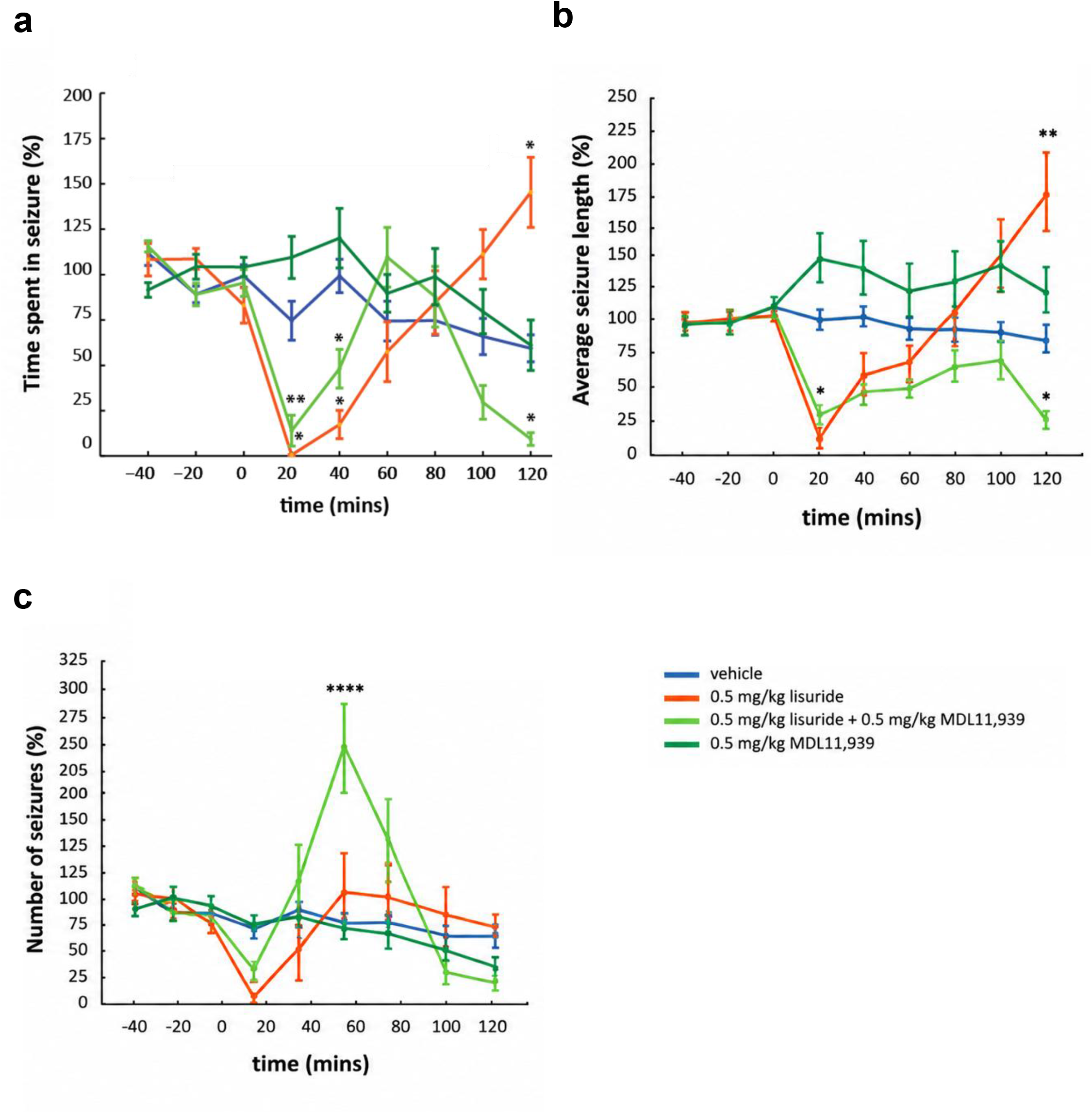
The 5-HT2A receptor antagonist MDL11,939 differentially modulates the biphasic effects of lisuride on ASs. **(a–c)** Time course of spontaneous absence seizure (AS) parameters following systemic administration of vehicle, lisuride (0.5 mg/kg), MDL11,939 (0.5 mg/kg), or lisuride (0.5 mg/kg) following 10 min pretreatment with MDL11,939 (0.5 mg/kg) (n=9 per group). **a**, time spent in seizures; **b**, mean seizure duration; c**C**, number of seizures. Pretreatment with MDL11,939 did not prevent the early anti-absence effect of lisuride but markedly modified its subsequent effects, preventing the late increases in time spent in seizures and seizure duration, and producing a transient increase in seizure number at 60 min. Data are expressed as a percentage of baseline and presented as mean ± SEM. Asterisks indicate significant differences at the indicated time points: \**p* < 0.05, \*\**p* < 0.01, and \*\*\*\**p* < 0.0001. Full statistical results from the two-way ANOVA, including the main effects of treatment and time, their interaction, and Dunnett’s post hoc comparisons are provided in Suppl. Table 1B.

In contrast, 0.5 mg/kg i.p. lisuride produced a pronounced biphasic response (Fig. 1A-C). Time spent in seizure was reduced by 99.2% at 20 min and 82.2% at 40 min (both *p* < 0.05), followed by a significant 118.4% increase at 120 min (*p* < 0.05). Average seizure length showed a similar profile, decreasing by 85.1% at 20 min (*p* < 0.01) and increasing by 101.9% at 120 min (*p* < 0.01). Seizure number was not significantly altered at individual time points, indicating that the delayed increase in seizure burden was predominantly associated with prolonged ASs.

### 3.2 5-HT2A receptor blockade differentially modifies the early and late effects of lisuride

To examine the contribution of 5-HT2ARs to lisuride’s effects, GAERS received the selective 5-HT2AR antagonist MDL11,939 (0.5 mg/kg, i.p.) 10 min before lisuride (0.5 mg/kg, i.p.). The lisuride-only and vehicle groups were the same animals included in the dose-response analysis; separate MDL11,939-only and lisuride-plus-MDL11,939 groups were added (*n* = 9 animals per group). Two-way ANOVA revealed significant effects of treatment and time, as well as a treatment × time interaction, on spontaneous absence seizures (Suppl. Table 1B). MDL11,939 did not prevent the early anti-absence effect of lisuride. Following combined treatment, time spent in seizure was reduced by 82.2% at 20 min and 54.4% at 40 min (both *p* < 0.05), while average seizure length was reduced by 74.6% at 20 min (*p* < 0.05). Thus, substantial early seizure suppression persisted during 5-HT2AR blockade.

In contrast, MDL11,939 reversed the delayed response to lisuride. At 120 min, lisuride alone increased time spent in seizure by 127.0% (*p* < 0.001) and average seizure length by 95.0% (*p* < 0.01), whereas following MDL11,939 pretreatment these measures were reduced by 85.4% and 73.0%, respectively (both *p* < 0.05). Combined treatment also produced a pronounced transient increase in seizure number at 60 min (257.2%, *p* < 0.0001), with no significant changes at other time points. MDL11,939 alone did not significantly alter time spent in seizure or seizure number at any individual time point, although a significant overall effect on average seizure length was observed (*p* < 0.01). Notably, lisuride was associated with agitation and erratic locomotor activity, which persisted following MDL11,939 pretreatment.

## 4. Discussion

This study demonstrates that systemic lisuride produces a dose- and time-dependent biphasic modulation of spontaneous ASs in GAERS. While 0.1 mg/kg elicited modest pro-absence effects, 0.5 mg/kg caused profound early suppression of ASs, followed by facilitation during the second hour. The early effect involved reductions in both seizure number and duration, whereas the delayed increase in seizure burden predominantly reflected in prolonged ASs. MDL11,939 did not prevent the early suppression but markedly blocked the delayed response, suggesting that distinct receptor mechanisms underlie the two phases of lisuride action.

These findings contrast with an earlier report of little effect of lisuride at 0.25–0.5 mg/kg but near-complete suppression of spike-and-wave discharges at 1 mg/kg, a dose that was associated with animal deaths [20]. This apparent discrepancy may reflect differences in experimental design, including the lack of habituation and the shorter baseline and, particularly, post-treatment observation periods, as the facilitatory effect observed here emerged only during the second hour which was not assessed in the study of Warter et al. (1988). This interpretation of differences in the recording protocol is supported by the limited involvement of 5-HT2Rs in ASs reported in earlier work from the same group [21], whereas subsequent studies using the same experimental design as the present work demonstrated robust 5-HT2AR-mediated modulation of ASs [8, 19].

Biphasic actions are not unique to the effects of lisuride on seizures. Temporally opposing effects on locomotor activity, with initial suppression followed by stimulation, have also been reported and attributed to the complex serotonergic and dopaminergic pharmacology of lisuride [22, 23]. Such observations suggest that temporally opposing responses may represent a broader feature of lisuride pharmacology rather than a phenomenon restricted to epileptic thalamocortical oscillations. More recent evidence indicates that lisuride instead displays relatively low efficacy across 5-HT2AR signaling pathways, including Gq and β-arrestin signaling, and, as a low-efficacy partial agonist, can display antagonist-like properties in the presence of higher-efficacy agonists [14]. Low 5-HT2AR signaling efficacy, rather than biased agonism per se, may therefore better distinguish lisuride from classical psychedelics [14, 24].

The early lisuride anti-absence effect differs from that of TCB-2, which suppresses ASs through 5-HT2AR activation when injected both systemically and within the VB [8, 19]. In contrast, MDL11,939 did not prevent lisuride-induced early suppression, indicating that this effect is largely 5-HT2AR-independent.

Lisuride’s broad receptor profile may account for this difference, as it acts at multiple serotonergic and dopaminergic receptors [14]. Dopaminergic mechanisms are particularly relevant because dopamine suppresses SWDs [20], while D2-like receptor activation reduces the enhanced tonic GABAA inhibition of VB TC neurons in GAERS [7, 25]. Conversely, 5-HT1A activation facilitates SWDs (Jakus et al., 2003), suggesting that lisuride’s early effect may reflect the combined action of multiple receptor systems.

Serotonergic and dopaminergic systems are closely interconnected [26]. In GAERS, nigrostriatal dopaminergic degeneration prolongs ASs and alters serotonergic transmission [27], suggesting that lisuride’s biphasic effects may reflect a changing dopaminergic-serotonergic balance. However, a dopaminergic contribution to the early effect remains to be directly demonstrated. In this respect, a key finding is the dissociation between seizure number and duration. At 20 min, lisuride reduced both parameters, whereas the delayed increase in seizure burden at 120 min predominantly reflected prolonged SWDs without increased seizure number. This suggests differential modulation of mechanisms governing SWD initiation and maintenance/termination in the biphasic lisuride effect.

This distinction is consistent with current concepts of ASs generation. SWDs initiation involves cortical networks followed by recruitment of cortico-thalamo-cortical circuits [2, 28]. Recent optogenetic evidence further shows that VB thalamocortical neurons can both initiate and terminate ASs depending on the state of the network [29], supporting partially distinct mechanisms controlling ASs occurrence and maintenance/termination. These processes may be differentially modulated by dopamine and serotonin, both of which are potent modulators of thalamic function [30, 31]. Consistent with this interpretation, MDL11,939 was ineffective on the early reduction in seizure burden and duration but prevented the delayed lisuride-induced ASs prolongation, by transiently increasing seizure number. Thus, the 5-HT2AR-sensitive component may preferentially modulate ASs maintenance/termination, whereas other lisuride-sensitive mechanisms may contribute to ASs initiation.

The present findings are also consistent with the evolving understanding of lisuride pharmacology. Although lisuride was historically considered a non-hallucinogenic 5-HT2AR agonist whose properties were attributed to biased [32], more recent evidence indicates relatively low efficacy across 5-HT2AR signaling pathways [14, 24, 33]. Low efficacy at 5-HT2ARs, together with activity at other serotonergic and dopaminergic receptors, may better explain its complex pharmacological profile and distinguish it from TCB-2.

The delayed response to lisuride should not imply that 5-HT2AR activation is intrinsically pro-absence, since both systemic TCB-2 and thalamic 5-HT2AR activation suppress ASs [8, 19]. Rather, 5-HT2AR signaling may contribute to the network state underlying the delayed response, potentially reflecting lisuride’s low efficacy and time-dependent recruitment of different receptor populations [14]. Behavioral state may also contribute, as ASs are vigilance-dependent and lisuride induced early agitation and locomotor activation.

In conclusion, lisuride’s biphasic effects appear to involve distinct receptor and network mechanisms. The early anti-absence effect is largely 5-HT2AR-independent, reducing both SWD occurrence and duration, whereas the delayed response predominantly reflects SWD prolongation and involves a 5-HT2AR-sensitive component. These findings support the concept that SWD initiation and maintenance/termination are partially dissociable processes and highlight the complex actions of multi-receptor drugs such as lisuride.

## Author contribution

MV: Methodology, Formal analysis, Writing - original draft. VC and GDG: Conceptualization, Funding acquisition, Resources, Methodology, Supervision, Writing - original draft, Writing - review and editing.

## Acknowledgments

This work was supported by Epilepsy Research UK (grant P1202 to VC and GDG), the Wellcome Trust (grant 91882 to VC) and the Ester Floridia Neuroscience Research Foundation (grant 1502 to VC).

---

**Supplementary Table 1.**
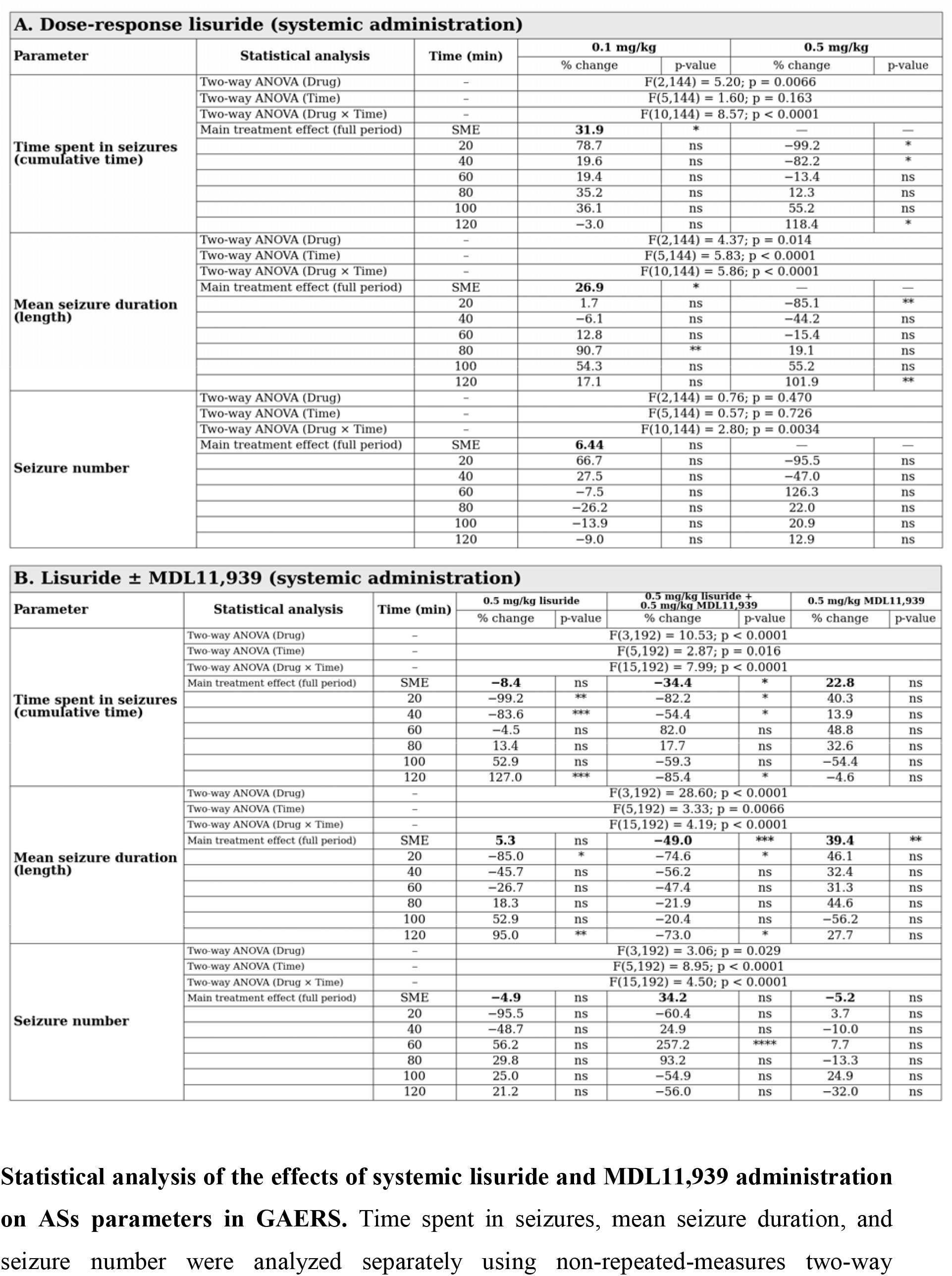
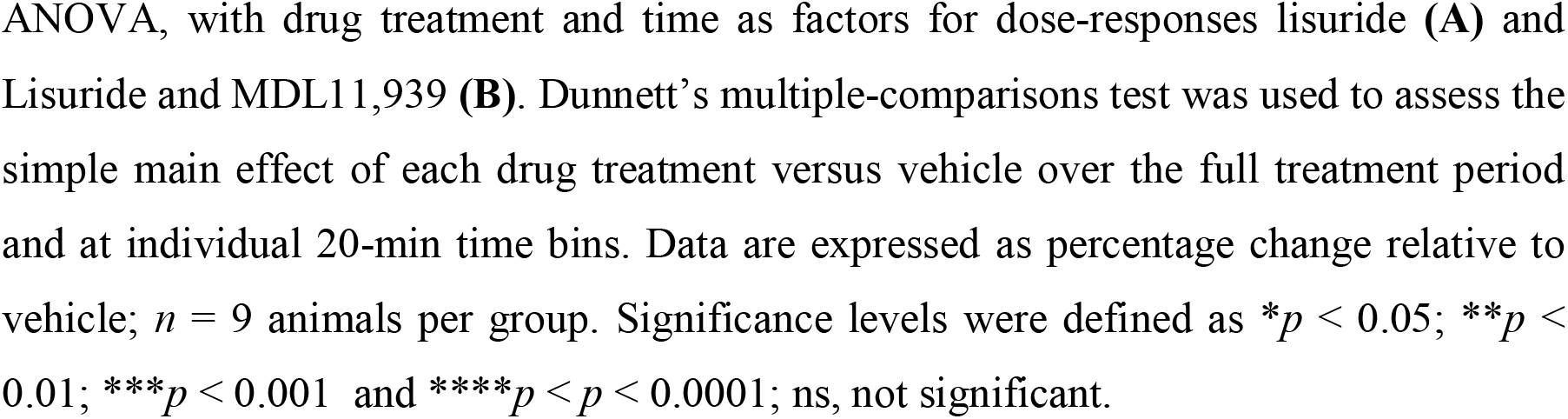
Summary statistics.

## References

1. Crunelli, V. and N. Leresche, Childhood absence epilepsy: genes, channels, neurons and networks. Nature Reviews Neuroscience, 2002. 3(5): p. 371–382.

2. Crunelli, V., et al., Clinical and experimental insight into pathophysiology, comorbidity and therapy of absence seizures. Brain, 2020. 143(8): p. 2341–2368.

3. Depaulis, A., O. David, and S. Charpier, The genetic absence epilepsy rat from Strasbourg as a model to decipher the neuronal and network mechanisms of generalized idiopathic epilepsies. J Neurosci Methods, 2016. 260: p. 159–74.

4. Barnes, N.M., et al., International Union of Basic and Clinical Pharmacology. CX. Classification of Receptors for 5-hydroxytryptamine; Pharmacology and Function. Pharmacological Reviews, 2021. 73(1): p. 310–520.

5. Cavaccini, A., et al., Thalamic distribution and effects of 5-HT2C receptors on tonic GABAA inhibition and absence seizures: implications for treatment. Frontiers in Pharmacology, 2026. Volume 17:1729460.

6. Crunelli, V. and G. Di Giovanni, Monoamine modulation of tonic GABAA inhibition. Rev Neurosci, 2014. 25(2): p. 195–206.

7. Cope, D.W., et al., Enhanced tonic GABAA inhibition in typical absence epilepsy. Nat Med, 2009. 15(12): p. 1392–8.

8. Morais, T.P., et al., Developmental and cell type-specific contributions of thalamic serotonin 2A receptors to absence seizures. Epilepsia, 2026. 67(8): p. 4305–4319.

9. Deidda, G., et al. The serotonin 2A receptor as a new therapeutic target to control absence seizures. in Mediterranean Neuroscience Society (MNS) Conference. 2019. Marrakech, Morocco.

10. Crunelli, V., et al., Thalamic 5-HT2A Receptor Activation Reduces GABA Uptake, Increases Tonic GABAA Inhibition and Induces Absence Seizures in Wistar Rats. Neuropharmacology, 2026: p. 111073.

11. Egan, C.T., et al., Agonist activity of LSD and lisuride at cloned 5HT2A and 5HT2C receptors. Psychopharmacology (Berl), 1998. 136(4): p. 409–14.

12. Cussac, D., et al., Agonist-directed trafficking of signalling at serotonin 5-HT2A, 5-HT2B and 5-HT2C-VSV receptors mediated Gq/11 activation and calcium mobilisation in CHO cells. Eur J Pharmacol, 2008. 594(1-3): p. 32–8.

13. Karaki, S., et al., Quantitative phosphoproteomics unravels biased phosphorylation of serotonin 2A receptor at Ser280 by hallucinogenic versus nonhallucinogenic agonists. Mol Cell Proteomics, 2014. 13(5): p. 1273–85.

14. Glatfelter, G.C., et al., Comparative Pharmacological Effects of Lisuride and Lysergic Acid Diethylamide Revisited. ACS Pharmacol Transl Sci, 2024. 7(3): p. 641–653.

15. Wallach, J., et al., Identification of 5-HT2A receptor signaling pathways associated with psychedelic potential. Nature Communications, 2023. 14(1): p. 8221.

16. Kimura, K., et al., Dual activation by lisuride of central serotonin 5-HT(1A) and dopamine D(2) receptor sites: drug discrimination and receptor binding studies. Behav Pharmacol, 1991. 2(2): p. 105–112.

17. Lidster, K., et al., Opportunities for improving animal welfare in rodent models of epilepsy and seizures. J Neurosci Methods, 2016. 260: p. 2–25.

18. Cope, D.W., S.W. Hughes, and V. Crunelli, GABAA receptor-mediated tonic inhibition in thalamic neurons. J Neurosci, 2005. 25(50): p. 11553–63.

19. Venzi, M., et al., Role for serotonin2A (5-HT2A) and 2C (5-HT2C) receptors in experimental absence seizures. Neuropharmacology, 2016. 108: p. 292–304.

20. Warter, J.-M., et al., Effects of drugs affecting dopaminergic neurotransmission in rats with spontaneous petit mal-like seizures. Neuropharmacology, 1988. 27(3): p. 269–274.

21. Marescaux, C., M. Vergnes, and A. Depaulis, Neurotransmission in rats’ spontaneous generalized nonconvulsive epilepsy. JOURNAL OF ANALYTICAL CHEMISTRY C/C OF ZHURNAL ANALITICHESKOI KHIMII, 1992: p. 335–335.

22. Adams, L.M. and M.A. Geyer, Patterns of exploration in rats distinguish lisuride from lysergic acid diethylamide. Pharmacology Biochemistry and Behavior, 1985. 23(3): p. 461–468.

23. Fink, H. and R. Morgenstern, Locomotor effects of lisuride: a consequence of dopaminergic and serotonergic actions. Psychopharmacology, 1985. 85(4): p. 464–468.

24. Ippolito, A., et al., Evidence that 5□HT2A receptor signalling efficacy and not biased agonism differentiates serotonergic psychedelic from non□psychedelic drugs. British Journal of Pharmacology, 2026. 183(14): p. 4058–4071.

25. Yagüe, J.G., et al., Dopaminergic modulation of tonic but not phasic GABAA-receptor-mediated current in the ventrobasal thalamus of Wistar and GAERS rats. Experimental neurology, 2013. 247: p. 1–7.

26. Di Giovanni, G., et al., Serotonin–dopamine interaction: electrophysiological evidence. Progress in brain research, 2008. 172: p. 45–71.

27. Tugba, E.K., et al., Prolongation of absence seizures and changes in serotonergic and dopaminergic neurotransmission by nigrostriatal pathway degeneration in genetic absence epilepsy rats. Pharmacology Biochemistry and Behavior, 2022. 213: p. 173317.

28. McCafferty, C., et al., Cortical drive and thalamic feed-forward inhibition control thalamic output synchrony during absence seizures. Nat Neurosci, 2018. 21(5): p. 744–756.

29. Morais, T.P., et al., Bidirectional control of generalized absence epilepsy networks via real-time direct depolarization of thalamocortical neurons. Prog Neurobiol, 2025. 252: p. 102774.

30. Monckton, J.E. and D.A. McCormick, Neuromodulatory role of serotonin in the ferret thalamus. J Neurophysiol, 2002. 87(4): p. 2124–36.

31. Govindaiah, G., Y. Wang, and C.L. Cox, Dopamine enhances the excitability of somatosensory thalamocortical neurons. Neuroscience, 2010. 170(4): p. 981–91.

32. González-Maeso, J., et al., Hallucinogens recruit specific cortical 5-HT(2A) receptor-mediated signaling pathways to affect behavior. Neuron, 2007. 53(3): p. 439–52.

33. Kehler, J. and M.S.T. Lindskov, Are the LSD-analogs lisuride and ergotamine examples of non-hallucinogenic serotonin 5-HT2A receptor agonists? J Psychopharmacol, 2025. 39(9): p. 889–895.

