## Supplementary Table 1 for "Biphasic Modulation of Absence Seizures by Lisuride in GAERS: Dissecting the Role of 5-HT2A Receptors"

**Running title**: Lisuride in Absence Seizures

### Keywords

Absence epilepsy; GAERS; Thalamocortical network; Ventrobasal thalamus; Spike–wave discharges; In vivo electrophysiology

**Supplementary Table 1. Summary statistics**

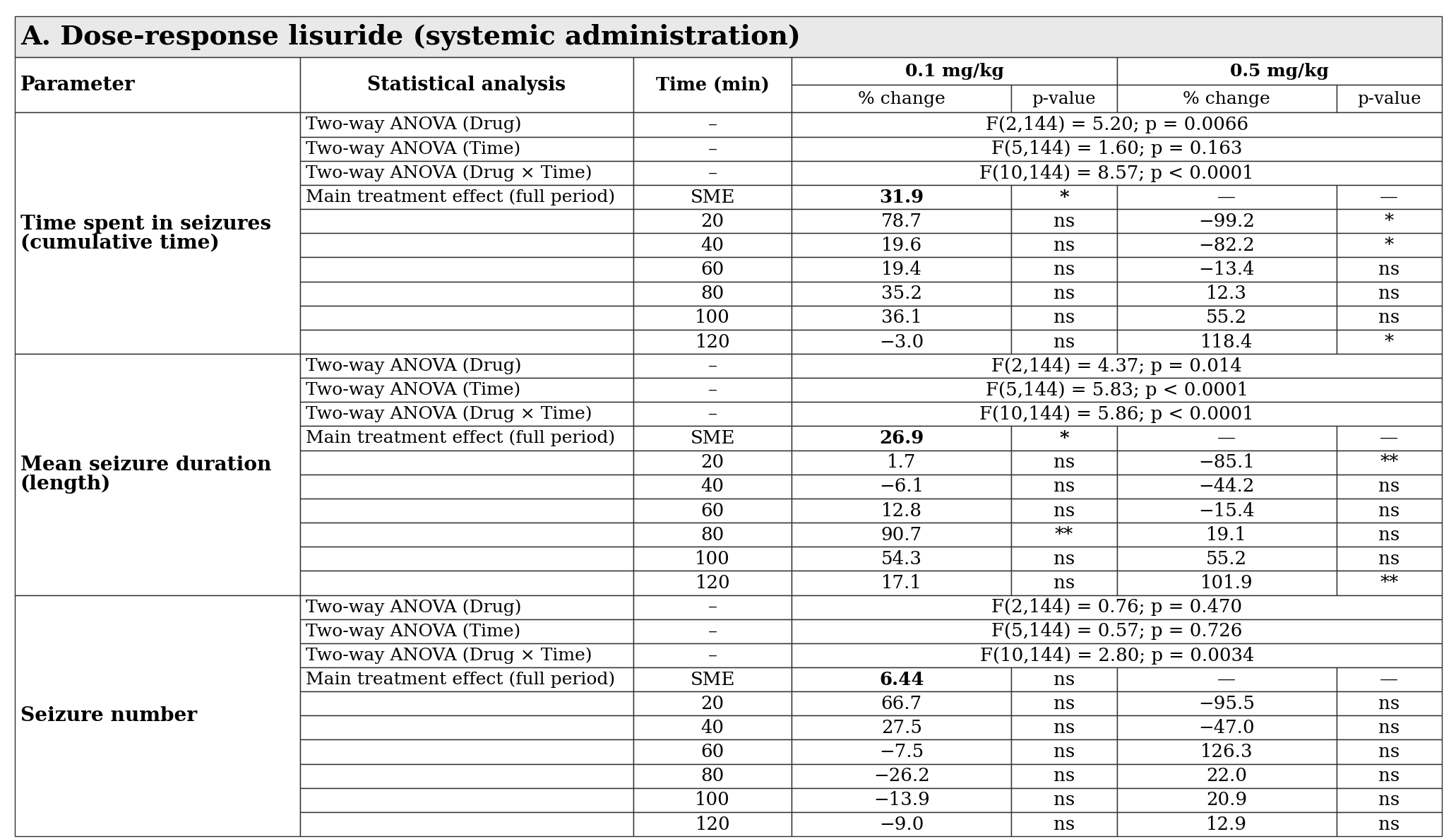

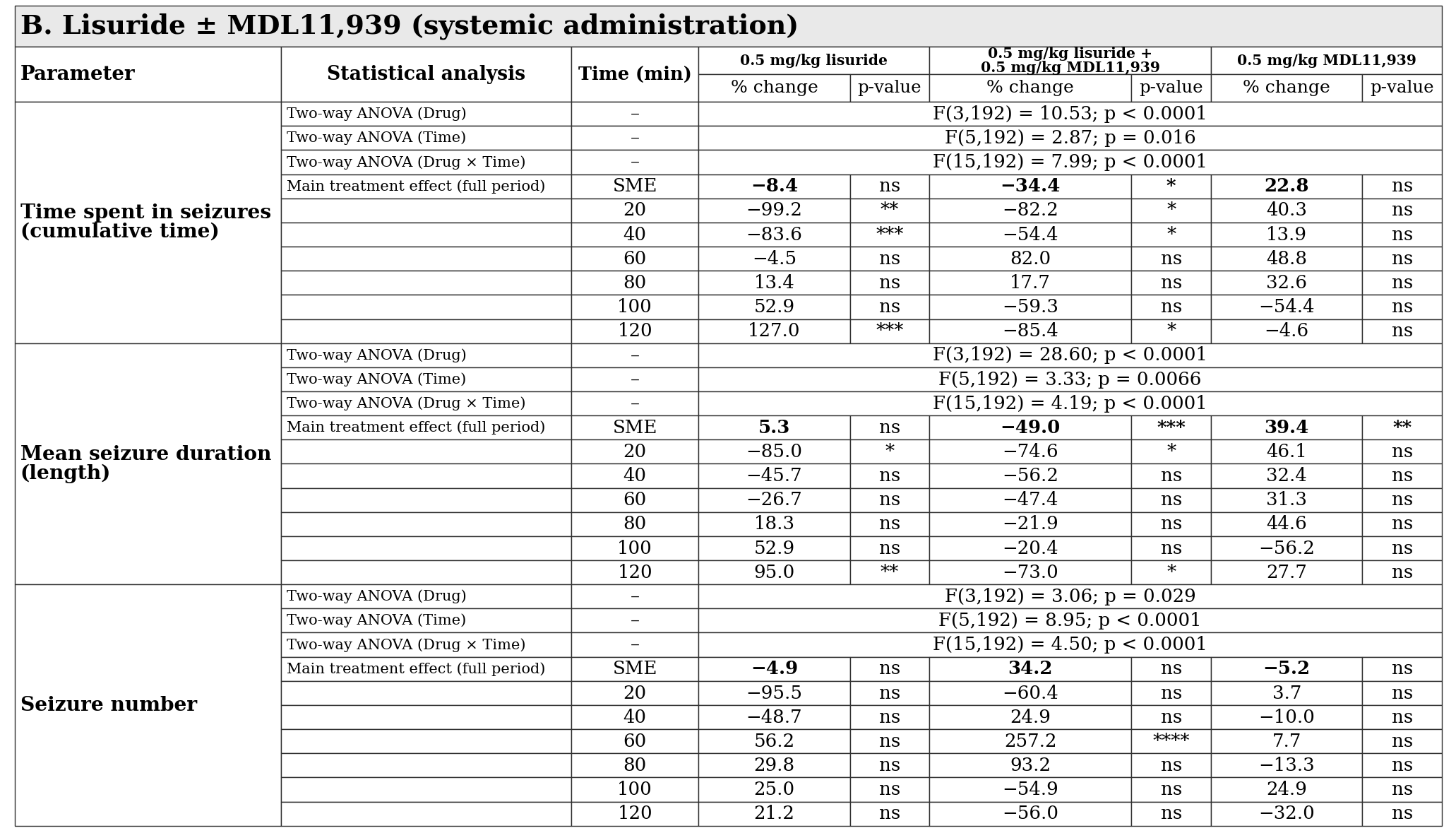

**Statistical analysis of the effects of systemic lisuride and MDL11,939 administration on ASs parameters in GAERS.** Time spent in seizures, mean seizure duration, and seizure number were analyzed separately using non-repeated-measures two-way ANOVA, with drug treatment and time as factors for dose-responses lisuride **(A)** and Lisuride and MDL11,939 **(B)**. Dunnett’s multiple-comparisons test was used to assess the simple main effect of each drug treatment versus vehicle over the full treatment period and at individual 20-min time bins. Data are expressed as percentage change relative to vehicle; n = 9 animals per group. Significance levels were defined as *p < 0.05; **p < 0.01; ***p < 0.001 and ****p < p < 0.0001; ns, not significant.
